# XY sex determination in a cnidarian

**DOI:** 10.1101/2022.03.22.485406

**Authors:** Ruoxu Chen, Steven M. Sanders, Zhiwei Ma, Justin Paschall, E. Sally Chang, Brooke M. Riscoe, Christine E. Schnitzler, Andreas D. Baxevanis, Matthew L. Nicotra

## Abstract

Sex determination occurs across animal species, but most of our knowledge about the mechanisms of sex determination comes from only a handful of bilaterian taxa. This limits our ability to infer the evolutionary history of sex determination within animals. In this study, we generated a linkage map of the genome of the colonial cnidarian *Hydractinia symbiolongicarpus* and used this map to determine that this species has an XX/XY sex determination system. We delineate the pseudoautosomal and non-recombining regions of the Y chromosome and show that the latter encodes a number of genes with male gonad-specific expression. These findings establish *Hydractinia* as a tractable non-bilaterian model system for the study of sex determination.

**Significance Statement:** What determines whether an animal is male or female? The answer depends on the species. Some rely on signals from their environment, while others take cues from their genome. Most of what we understand about sex determination comes from traditional model organisms (such as mice, flies, and worms) or groups of well-studied vertebrates and insects. Studying sex determination in other animals, especially those from phyla that diverged early in animal evolution, would allow us to better understand how sex has evolved across the animal kingdom and could reveal pathways present in the eumetazoan ancestor >600 million years ago. In this study, we show that the cnidarian *Hydractinia symbiolongicarpus* has XY sex determination, establishing this species as a model system for sex determination in an understudied group of animals.

## INTRODUCTION

Sex determination governs whether a gonad develops into an ovary or testis (Capel 2017). In animals, the primary sex determination signal can be genetic or environmental (Picard et al. 2021). Genetic sex determination systems include sex chromosomes with male (XY) or female (ZW) heterogamety, systems in which males are haploid and females are diploid, or systems in which sex is determined by dosage at one or more autosomal loci. Environmental sex determination systems are similarly diverse, relying on external cues such as temperature, photoperiod, food, or social environment. In either case, the primary sex determination signal typically resides atop a cascade of pathways leading to the differentiation of either male or female gonads (Adolfi et al. 2021). The fact that many genes from these pathways are conserved across animals has inspired hypotheses about how sex determination mechanisms evolve (Wilkins 1995; Graham et al. 2003; Herpin and Schartl 2015) and has also led to speculation about the nature of ancestral sex determination systems. This latter issue is difficult to address because most of what we know about animal sex determination comes from vertebrates, arthropods, nematodes, and a handful of other bilaterians (Kato et al. 2011; Chong et al. 2013; The Tree of Sex Consortium 2014; Picard et al. 2021). Therefore, it is important to investigate sex determination across a broader diversity of species, especially nonbilaterians.

Cnidarian sex determination systems would be particularly informative in this context, because the phylum is the sister group to all bilaterians (Whelan et al. 2017; Simion et al. 2017). Cnidarians exhibit a range of sexual strategies, including gonochorism (separate sexes), simultaneous hermaphroditism, and sequential hermaphroditism (Siebert and Juliano 2017). Although sexual differentiation and gametogenesis are well-studied in several cnidarian species (Siebert and Juliano 2017), the primary sex determination signal has not been identified in any cnidarian. In fact, the only data directly addressing sex determination come from studies in two coral species. In the first, a karyotype of *Acropora solitaryensis* was characterized via fluorescence *in situ* hybridization with DNA extracted from sperm and eggs, revealing a putative Y chromosome (Taguchi et al. 2014). In the second, a genome-wide analysis of SNPs from field-collected red corals (*Corallia rubrum*) yielded several male-specific loci consistent with chromosomal XX/XY sex determination (Pratlong et al. 2017).

The hydroid *Hydractinia symbiolongicarpus* is a promising laboratory model system for cnidarian sex determination. *Hydractinia* colonies are gonochoristic and release gametes daily in response to a light cue. Fertilization is external, and each embryo develops into a crawling larva that metamorphoses into a primary feeding polyp. This polyp extends stolons from its base from which additional feeding polyps (gastrozooids) grow, thus creating a multi-polyp colony (**Fig. 1A**). Within 3-4 months, colonies develop polyps specialized for reproduction (gonozooids) in which eggs or sperm are easily observed (**Fig. 1B,C**). In laboratory settings, experimental crosses produce offspring with a 1:1 sex ratio (Hauenschild 1954). Although anecdotal evidence from laboratory crosses suggest the sex ratio is not affected by environmental conditions (Matthew Nicotra, unpublished data), the primary sex determination signal is unknown. Most importantly, *Hydractinia* is a tractable genetic model system. It is easily bred in the laboratory, has a sequenced genome, and is amenable to gene knockdown, knockout, and knockin (Frank et al. 2020).

**Fig. 1.**
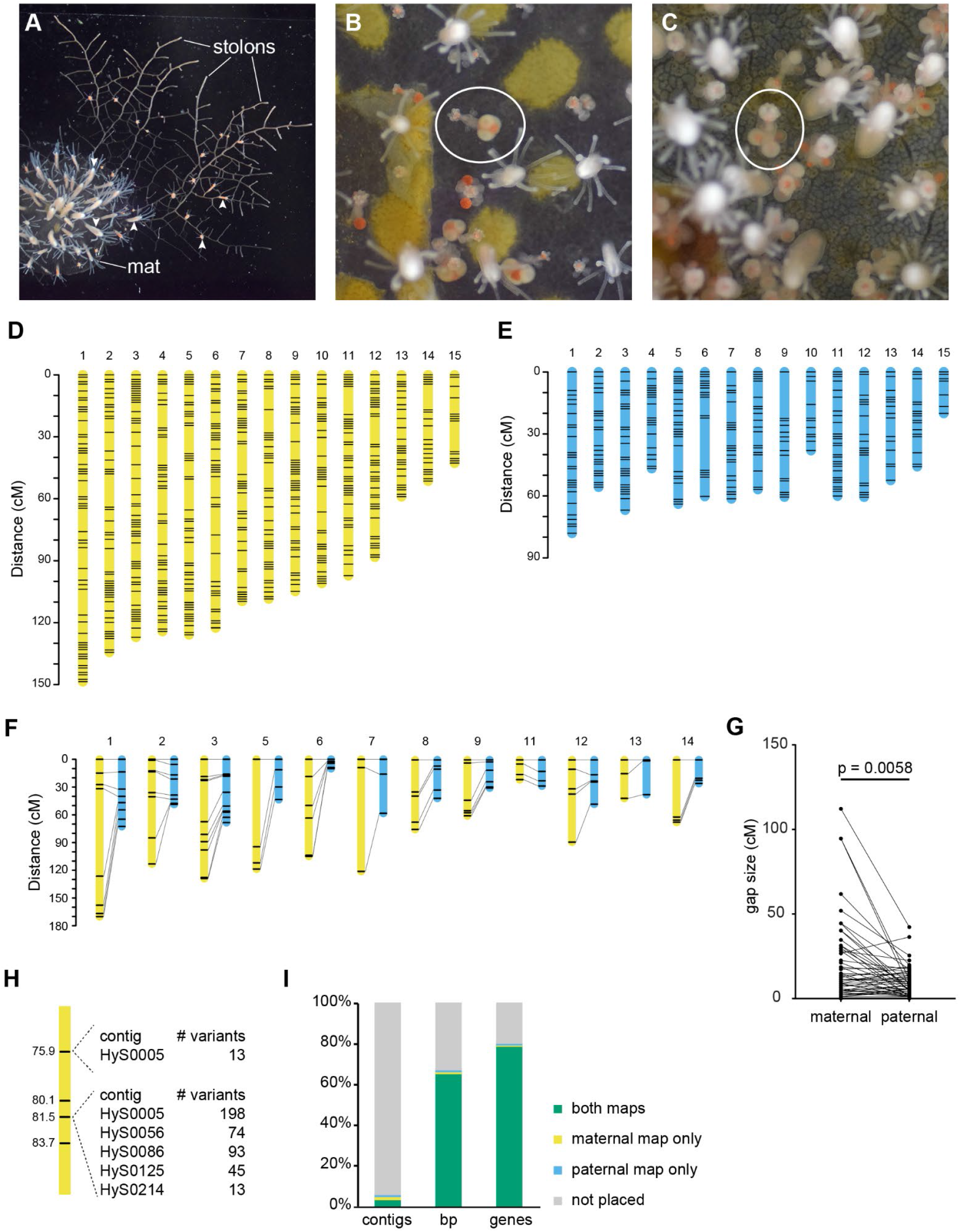
*Hydractinia* colonies and linkage maps. *(A)* An immature *Hydractinia* colony. Arrowheads indicate gastrozooids. *(legend continues on next page)* *(B)* Female gonozooid (circled) with one mature gonophore containing mature eggs. *(C)* Male gonozooid (circled). *(D)* Maternal linkage map. *(E)* Paternal linkage map. *(F)* Comparison of markers at equivalent physical locations in maternal and paternal maps. Lines connect markers located within 5 kb of each other in the reference genome. *(G)* Comparison of gap sizes in maternal and paternal linkage maps. Gap sizes from equivalent markers are connected by lines. *(H)* Example of how markers in the linkage map are bins of multiple variants. This example shows four markers from maternal linkage group 1. The marker at 75.9 cM represents 13 variants from one contig. The marker at 81.5 cM represents 423 variants from five contigs. *(I)* Representation of the *Hydractinia* genome assembly in the maternal and paternal linkage maps. Results are shown as a percent of the total number of contigs, base pairs (bp), or annotated genes.

In *Hydractinia*, sexual differentiation and gametogenesis are processes that occur continuously throughout the colony’s life within each gonozooid (Weismann 1883; Müller 1964). *Hydractinia* stem cells, known as i-cells, migrate into gonozooids, where they acquire germ cell fate and become gamete progenitors (DuBuc et al. 2020). These progenitors then migrate from the neck of the gonozooid into sporosacs, where they mature into either eggs or sperm. Although it is unclear when sex determination takes place during this process, it is clear that the sexual identity of the colony is dependent on identity of the i-cell. Evidence for this comes from experiments in which a donor colony is grafted to an i-cell-depleted recipient that then begins to produce gametes matching the sex of the donor (Müller 1964, 1967; Muller et al. 2004).

Although the sex of a colony is usually stable for its entire life, there have been reports of colonies that produce male and female gametes simultaneously (Bunting 1894; Hauenschild 1954; Mali et al. 2011). In each case, the colony produces gonozooids with sporosacs containing mature sperm and immature eggs. In two cases, these colonies were reared from a single primary polyp, thus ruling out the possibility that male and female colonies had fused together and enabled i-cells of both sex to enter the same gonozooid (Hauenschild 1954; Mali et al. 2011). In our own laboratory, we have observed rare colonies that reach sexual maturity as fully functional females and then begin to produce male gametes that appear to overtake the entire colony (Leo Buss & Matthew Nicotra, unpublished observations). In these cases, it has been unclear whether the intersex colony is the product of an unnoticed fusion between males and females.

In this study, we sought to test the hypothesis that *Hydractinia* has genetic sex determination. To that end, we constructed a genome-wide linkage map and used it to identify sex-linked loci. We find a pattern consistent with XY sex determination. The Y chromosome is estimated to have half non-recombining region and half pseudoautosomal regions. The non-recombining region spans at least 7 Mb and consists of at least 461 annotated genes. Within this pool of candidate genes, several are expressed exclusively in male sexual polyps, making them prime candidates for further study.

## RESULTS

### A linkage map of the *Hydractinia* genome

To create a *Hydractinia* linkage map, we generated a population of 90 F_1_ offspring by breeding a male colony (291-10) to a half-sibling female colony (295-8; *SI Appendix*, Fig. S1). Offspring and parents were then sequenced using Illumina sequencing methods to yield a mean coverage of 47 ± 11x per sample, with a mean depth of 162 ± 38 million reads per sample. (For per-sample statistics, see detailed methods in *SI Appendix* and Dataset S1.) Reads were mapped to a reference assembly of the paternal genome, and a total of 9.74 million single nucleotide polymorphisms (SNPs) and 1.83 million insertion/deletion variants (indels) were identified. After filtering low quality variant calls, 1,312,632 variants (1,083,937 SNPs and 228,695 indels) remained.

From this filtered dataset, we identified markers suitable for linkage mapping via a pseudo-testcross strategy (Grattapaglia and Sederoff 1994). In a pseudo-testcross, parents from an outcrossing population are bred to create an F_1_ population. A genetic map for each parental genome is then constructed from markers that are heterozygous in that parent and homozygous in the other. For example, a genetic map of the maternal genome can be constructed from variants that are heterozygous in the female parent and homozygous in the male parent. Likewise, a genetic map of the paternal genome can be constructed from variants heterozygous in the male parent and homozygous in the female parent. In this study, we identified 303,020 variants (255,004 SNPs and 48,016 indels) suitable for mapping the maternal genome and 251,912 variants (217,242 SNPs and 34,670 indels) suitable for mapping the paternal genome. SNPs and indels were treated equivalently for mapping purposes. After removing variants that displayed segregation distortion at *p* < 10^−5^ and thinning the remaining variants such that the minimum distance between adjacent variants was 5,000 bp, we were left with 23,462 variants (20,058 SNPs and 3,404 indels) for the maternal genome and 22,359 variants (19,771 SNPs and 2,863 indels) for the paternal genome (Datasets S2 and S3).

We constructed linkage maps in R (version 3.6.1; (R Core Team 2019) with the package OneMap (Version 2.1.1) (Margarido et al. 2007). Variants with identical genotypes in the F_1_ animals were binned to create single markers for linkage mapping. After binning, we recalculated segregation distortion using a Bonferroni-corrected *p*-value of 0.05 and removed any remaining distorted markers. This resulted in a set of 977 markers for the maternal genome and 487 for the paternal genome. Markers were placed into linkage groups with a maximum recombination frequency of 0.4 and minimum LOD score of 6.14 and 5.56 for the female and male datasets, respectively. In both cases, 15 linkage groups were obtained. This matched the predicted haploid chromosome number (n = 15) obtained from a separate karyotype analysis (*SI Appendix*, Fig S2).

When constructing linkage maps, OneMap often placed two markers within <0.0001 cM of each other. On closer inspection, we found these markers always represented variants with an identical segregation pattern in the F_1_ population, except that they were in the opposite phase. This artificially inflated the number of informative markers in each linkage map. Therefore, we removed one marker from each pair, except in cases where the markers indeed represented different contigs (see detailed experimental methods in *SI Appendix*). We then removed any remaining markers that could not be confidently placed, resulting in the final maps.

The final map of the maternal genome consisted of 590 markers and spanned 1545.5 cM (**Fig. 1D** and *SI Appendix*, Fig. S3, Dataset S4), with an average gap of 2.69 cM/marker. The final map of the paternal genome consisted of 305 markers spanning 827.7 cM (**Fig. 1E** and *SI Appendix*, Fig. S4, Dataset S5), with an average gap of 2.85 cM/marker. Synteny between the maternal and paternal maps was determined by identifying linkage groups with a preponderance of markers from the same genomic contigs. Linkage groups were numbered according to decreasing genetic (cM) length in the maternal map.

We immediately noticed that the maternal linkage map was nearly twice as long as the paternal map, suggesting the genome-wide recombination rate was higher in the female parent compared to the male. Alternatively, the maternal map might be longer because it incorporated more markers and was more likely to be inflated by genotyping errors (Ronin et al. 2017). To test these hypotheses, we compared rates of recombination between equivalent physical locations in the maternal and paternal genomes. To do this, we first identified pairs of markers – one maternal and one paternal – located within 5 kb of each other in the reference genome. We then used these markers to reconstruct each linkage group, allowing us to directly compare rates of recombination between nearly identical locations in the maternal and paternal genomes. Of the 12 reconstructed linkage groups, 11 were longer in the maternal map (**Fig. 1F**). Gaps in the maternal map were also larger than the equivalent gaps in the paternal map (*p* = 0.0058, Wilcoxon matched-pairs signed rank test; **Fig. 1G**) and the average gap size was larger in the maternal map (20.2 cM) than the paternal map (9.3 cM). These data were consistent with a higher recombination rate in the female genome.

We next sought to determine how much of the *Hydractinia* genome assembly could be placed on each linkage map. Since each marker was a bin of variants from the reference genome, many markers represented variants from several contigs (**Fig. 1H**). To account for this, we created detailed linkage maps in which each marker was ‘unbinned’ and each underlying variant assigned a genetic position (Datasets S6 and S7). We then determined how many contigs could be placed on each map. In all, 273 contigs, representing 67.6% of the 403,824,369 bp assembly and nearly 81% of the 22,022 annotated genes could be placed in at least one linkage map (**Fig. 1I**, *SI Appendix*, Table S1). This analysis also allowed us to identify 15 contigs that were split between different linkage groups and may represent misassemblies in the reference genome (*SI Appendix*, Table S2).

### *Hydractinia* has an XY sex determination system

We recorded the sex of each F_1_ animal as soon as it could be determined from its developing gonophores (**Fig. 2A-B**). We then monitored the animals biweekly to identify instances of sexual chimerism. Most colonies (87/90) remained a single sex for the entire study, with an overall sex ratio that did not differ significantly from 1:1 (40:47 male:female; chi squared goodness of fit test χ^2^ = 0.563, *p* = 0.453). In contrast, three male colonies appeared to change sex. Two began to develop female gonophores approximately six months after being classified as male (**Fig. 2C and D**). For both colonies, we explanted fragments bearing only male or female gonozooids to new slides and monitored them. These new colonies have remained either male or female for >37 months. A third animal (339-083) was also initially classified as male but, six months later, was found to have only female gonozooids. While this animal may have undergone a complete sex change between our biweekly observations, we were unable to rule out the possibility that it we had mislabeled it as male in our records, and this error had gone unnoticed for several months. We therefore excluded it from subsequent analyses.

**Fig. 2.**
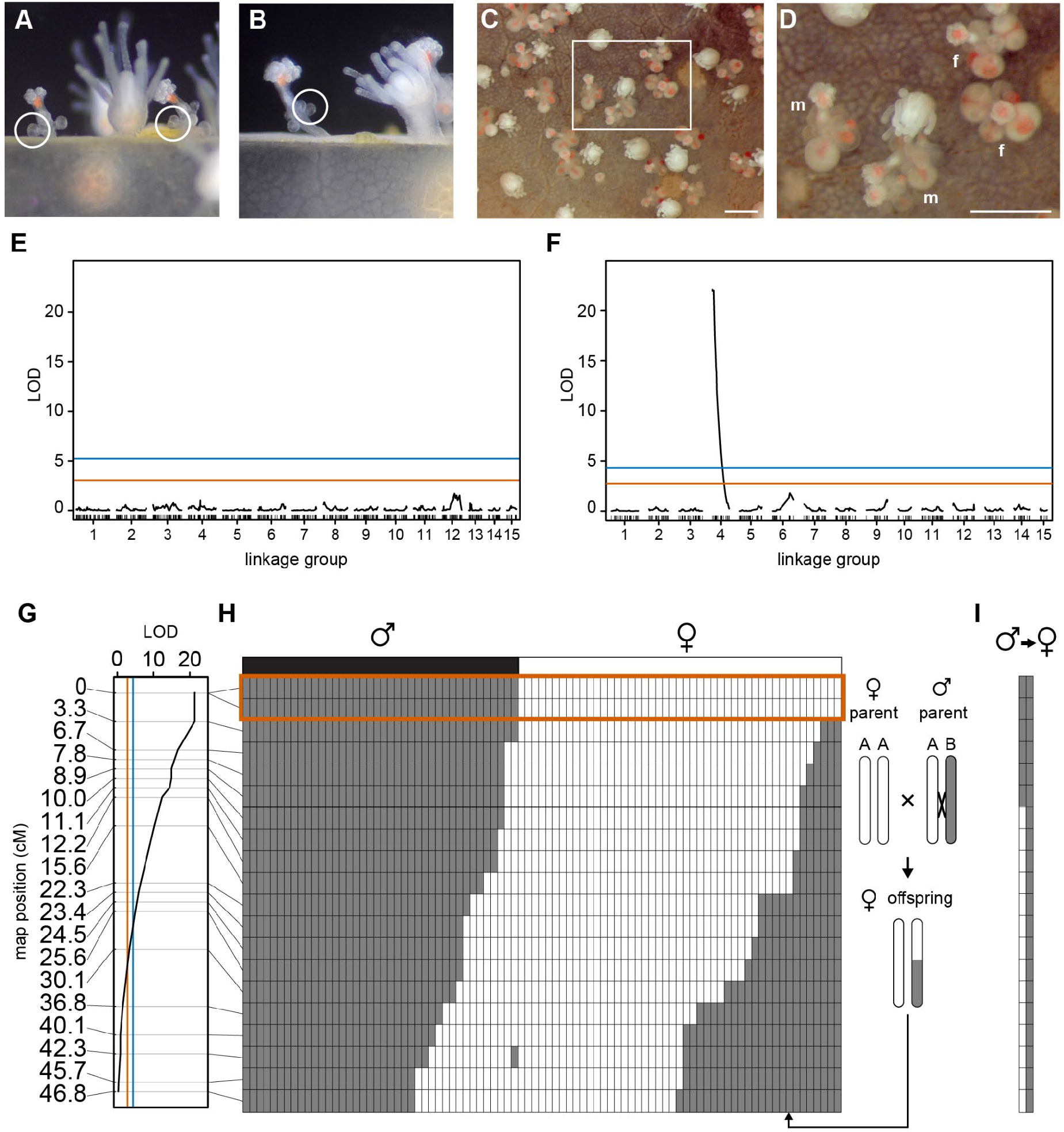
XY sex determination in *Hydractinia*. *(A)* Early and late stages of development of female gonozooids *(B)* Early and late stages of development of male gonozooids *(C)* Image of colony 339-116 on the day the first female gonophores were identified. *(D)* Close-up of boxed area in (C). m, gonozooid bearing male gonophores. f, gonozooid bearing female gonophores. Scale bars in C and D are 1 mm. *(E)* LOD chart of QTL for sex in the maternal linkage map. *(G)* Detail of LOD chart for linkage group 4 from the paternal linkage map. In *E-G*, significance thresholds of *p =* 0.05 and *p =* 10^−5^ are denoted in red and blue, respectively. *(H)* Recombination map of linkage group 4. Plot depicts genotype segregation pattern of pseudo-testcross markers (rows) in the F_1_ progeny (columns). Genotypes: AA: white; AB, gray. An example of a recombinant progeny is illustrated to the right of the plot. Observed sex of each F_1_ progeny is displayed above the plot. Genotypes in red box show perfect correlation with sex phenotype. *(I)* Recombination map of linkage group 4 in two animals with sexual chimerism.

We next used R/qtl (Broman et al. 2003) to identify markers associated with the initial sex of each animal. The analysis was performed independently on the maternal and paternal maps. While no significant loci were identified in the maternal map (**Fig. 2E**), several loci with a strong association with sex were located on the paternal map (**Fig. 2F**). The markers with the most significant association (p << 10^−5^) were located at 0 and 3 cM on linkage group 4 (**Fig. 2G**). Excluding the two sexual chimeras from this analysis did not affect these results significantly (*SI Appendix*, Fig. S5).

Next, we sought to determine how well the markers on linkage group 4 correlated with the intial sex of the colony. Using the linkage phase information estimated by OneMap, we determined the genotype of each F_1_ animal at each marker, which allowed us to infer which paternal haplotype had been inherited and whether it was a product of recombination (**Fig. 2H**). For animals with a stable sex, we found a perfect correlation between their sex and their genotype at the two 0 cM markers (**Fig. 2H**, red box). The two male to female sexual chimeras also carried a ‘male’ genotype at these markers (**Fig. 2I**). These data, combined with the fact that sex-linked markers were only identified on the male map, indicate *Hydractinia* has an XY sex determination system, and that the sex locus is located at the end of linkage group 4.

### The *Hydractinia* Y chromosome has a large pseudoautosomal region

As sex chromosomes evolve from autosomes, the sex-limited chromosome (Y or W) typically develops recombination suppression followed by degeneration and gene loss in the non-recombining region (Ponnikas et al. 2018). This non-recombining region is often linked to a pseudoautosomal region that continues to recombine with its heterologous counterpart (X or Z). To determine the extent of these regions on the *Hydractinia* Y chromosome, we drew connections between the maps of linkage group 4 and the reference genome assembly (**Fig. 3**). To simplify the visualization, each marker was mapped to the physical position of one representative variant per contig. Six contigs from the reference assembly could only be connected to one of the linkage maps. These contigs may represent sequences that have significantly diverged or been lost on either the X or the Y chromosome. Alternatively, they may not have been placed in both maps because they did not possess heterozygous variants in one parent.

**Fig. 3.**
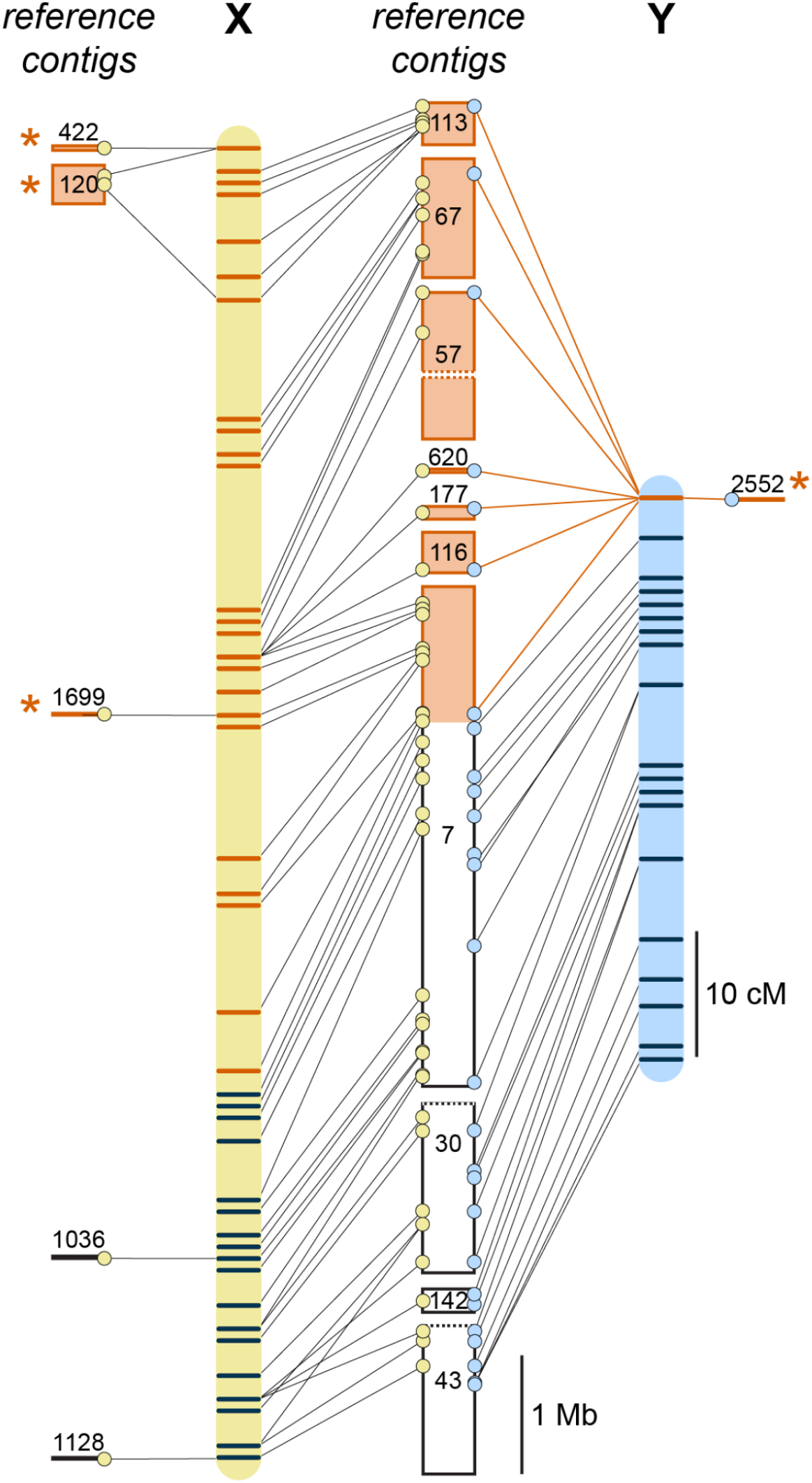
Genomic positions of markers in the maternal (yellow) and paternal (blue) maps of linkage group 4. Regions and markers outlined in orange are located within the non-recombining regions of the X and Y chromosome. Markers indicated in black are located in the pseudoautosomal region.

On the paternal map, which represents recombination between the X and Y chromosomes, the two 0 cM markers corresponded to a non-recombining region spanning at least 6.46 Mbp. This same genomic region corresponded to 87.6 cM on the maternal linkage map, indicating ample recombination between X chromosomes in the maternal genome. The pseudoautosomal region spanned at least 10.6 Mbp and corresponded to 36.7 cM in the maternal map and 46.8 cM in the paternal map. Together, these data indicate that more than half of the *Hydractinia* Y chromosome is a pseudoautosomal region.

### Gene content of the *Hydractinia* sex locus

The genomic contigs that align with the non-recombining regions of the X and Y chromosomes contain 461 annotated genes (*SI Appendix*, Dataset S8). Since our genome assembly is a haploid representation of the diploid paternal genome, we cannot assign these genes to specific sex chromosomes. Annotation of these genes revealed a variety of predicted functions (*SI Appendix*, Dataset S9) but no obvious homologs of genes or domains found in other animal sex determination pathways.

Sex-specific differences in gene expression can be a useful metric for identifying candidate sex determination genes. Using previously published data (Sanders et al. 2014), we calculated the average expression of each gene in gastrozooids, male gonozooids, and female gonozooids. A reasonable assumption for a sex determination gene is that it is only expressed in gonozooids. When we set a threshold of >1 fragments per kilobase per million mapped reads (FPKM) for a gene to be considered ‘expressed’, we found 25 genes with gonozooid-only expression. Of these, 18 were expressed only in male gonozooids, four were expressed only in female gonozooids, and three were expressed in male and female gonozooids (Table 1 and *SI Appendix*, Dataset S9). In the course of our analysis, we noticed several genes with high expression in either male or female gonozooids but low expression in gastrozooids. Therefore, we ran a second calculation using 5 FPKM as the expression threshold. At this level, we identified 16 additional gonozooid-specific genes, of which 13 were male-specific and three were female-specific (Table 1). In both analyses, there were more male gonozooid-specific than female gonozooid-specific genes and, moreover, the male-specific genes were more highly expressed than the female-specific ones.

**Table 1.** Genes with gonozooid- and sex-specific expression in the non-recombining region of the Hydractinia X and Y chromosomes.

| Gene ID | Expression (FPKM) |  |  | Top BLASTP hit (vs nr) | E-value |
| --- | --- | --- | --- | --- | --- |
|  | gastro | female gono | male gono |  |  |
| Expression threshold = 1 |  |  |  |  |  |
| HyS0057.51 | 0.84 | 0.03 | 314.84 | neural Wiskott-Aldrich syndrome protein-like (*Dendronephthya gigantea*) | 7.00E-08 |
| HyS0057.46 | 1.00 | 0.00 | 192.60 | potassium/sodium hyperpolarization-activated cyclic nucleotide-gated channel 1-like (*Hydra vulgaris*) | 3.50E-12 |
| HyS0057.47 | 0.00 | 0.00 | 67.81 | potassium/sodium hyperpolarization-activated cyclic nucleotide-gated channel 1-like (*Hydra vulgaris*) | 2.60E-35 |
| HyS0067.10 | 0.16 | 0.00 | 30.26 | - |  |
| HyS0113.16 | 0.76 | 0.00 | 6.41 | hypothetical protein BSL78_23407 (*Apostichopus japonicus*) | 6.80E-22 |
| HyS0067.3 | 0.45 | 0.03 | 5.67 | uncharacterized protein (*Exaiptasia pallida*) | 5.30E-56 |
| HyS0113.15 | 0.00 | 0.00 | 4.61 | uncharacterized protein (*Hydra vulgaris*) | 5.50E-34 |
| HyS0067.73 | 0.00 | 0.00 | 4.37 | uncharacterized protein (*Astyanax mexicanus*) | 3.80E-11 |
| HyS0007.328 | 0.27 | 0.24 | 3.76 | - |  |
| HyS0057.18 | 0.00 | 0.00 | 3.30 | hypothetical protein (*Exaiptasia pallida*) | 6.30E-10 |
| HyS0067.71 | 0.59 | 0.00 | 2.85 | uncharacterized protein (*Orbicella faveolata*) | 7.60E-07 |
| HyS0057.124 | 0.00 | 0.09 | 1.81 | COUP-TF protein, partial (*Hydractinia echinata*) | 3.80E-72 |
| HyS0057.129 | 0.31 | 0.12 | 1.82 | COUP-TF protein, partial (*Hydractinia echinata*) | 2.70E-175 |
| HyS0116.7 | 0.93 | 0.91 | 2.20 | uncharacterized protein (*Orbicella faveolata*) | 2.50E-60 |
| HyS0057.71 | 0.67 | 0.06 | 1.20 | hypothetical protein (*Trichomalopsis sarcophagae*) | 4.20E-43 |
| HyS0067.4 | 0.06 | 0.00 | 1.07 | Methylmalonic aciduria and homocystinuria type D-like, mitochondrial (*Exaiptasia pallida*) | 6.10E-05 |
| HyS0007.327 | 0.00 | 0.00 | 1.02 | - |  |
| HyS0057.113 | 0.44 | 0.51 | 1.27 | putative SCAN domain-containing protein 3-like (*Apostichopus japonicus*) | 2.30E-37 |
| HyS0007.332 | 0.36 | 9.66 | 0.00 | - |  |
| HyS0116.3 | 0.00 | 5.77 | 0.00 | uncharacterized protein (*Dendronephthya gigantea*) | 7.80E-22 |
| HyS0120.5 | 0.00 | 1.64 | 0.00 | - |  |
| HyS0007.350 | 0.00 | 1.35 | 0.87 | - |  |
| Expression threshold = 5 |  |  |  |  |  |
| HyS0007.412 | 2.83 | 0.40 | 943.95 | uncharacterized protein (*Hydra vulgaris*) | 3.10E-05 |
| HyS0057.61 | 2.08 | 0.00 | 859.64 | E3 ubiquitin-protein ligase MARCH2-like (*Hydra vulgaris*) | 2.70E-63 |
| HyS0057.48 | 2.84 | 0.04 | 531.51 | Potassium/sodium hyperpolarization-activated cyclic nucleotide-gated channel 2, partial (*Charadrius vociferus*) | 1.00E-60 |
| HyS0057.36 | 4.26 | 0.27 | 475.83 | - |  |
| HyS0057.16 | 3.91 | 3.05 | 32.44 | protein FAM221B-like (*Dendronephthya gigantea*) | 2.40E-91 |
| HyS0113.24 | 4.63 | 0.06 | 12.41 | hypothetical protein M513_02985, partial (*Trichuris suis*) | 1.50E-15 |
| HyS0057.114 | 2.15 | 0.21 | 11.43 | uncharacterized protein LOC109593289 (*Amphimedon queenslandica*) | 3.80E-10 |
| HyS0120.42 | 2.88 | 0.35 | 9.24 | mitogen-activated protein kinase kinase kinase 12-like (*Hydra vulgaris*) | 1.00E-27 |
| HyS0007.368 | 3.36 | 0.24 | 8.58 | A disintegrin and metalloproteinase with thrombospondin motifs 6-like (*Stylophora pistillata*) | 1.70E-79 |
| HyS0067.49 | 0.80 | 2.18 | 10.50 | uncharacterized protein (*Hydra vulgaris*) | 8.70E-140 |
| HyS0067.65 | 1.39 | 0.27 | 7.65 | alpha-1-macroglobulin-like (*Hydra vulgaris*) | 3.40E-198 |
| HyS0057.88 | 2.81 | 0.19 | 6.35 | uncharacterized protein (*Hydra vulgaris*) | 6.30E-24 |
| HyS1699.2 | 4.16 | 2.40 | 5.84 | fanconi-associated nuclease 1-like (*Hydra vulgaris*) | 3.00E-92 |
| HyS0007.387 | 2.71 | 3.01 | 5.09 | DNA topoisomerase 3-alpha-like (*Hydra vulgaris*) | 2.60E-246 |
| HyS0007.322 | 3.09 | 8.03 | 4.23 | importin-11 (*Hydra vulgaris*) | 2.20E-259 |
| HyS0067.52 | 2.85 | 5.66 | 1.03 | fibroblast growth factor receptor-like (*Hydra vulgaris*) | 7.80E-110 |
| HyS0067.50 | 1.91 | 5.21 | 0.58 | fibroblast growth factor receptor-like (*Hydra vulgaris*) | 6.00E-110 |

## DISCUSSION

In this study, we have demonstrated that *H. symbiolongicarpus* has an XX/XY genetic sex determination system. We also characterized the non-recombining region of the Y chromosome, which spans at least 6.46 Mbp. This region contains at least 461 genes, many of which are expressed only in male gonozooids and are good candidates for sex determination genes.

Although *Hydractinia* has sex chromosomes, how they determine sex remains unclear. Sex determination via X and Y chromsomes can occur in several ways. In mammals, for instance, the Y chromosome encodes the *sex determining region Y* (*Sry*) gene, a transcription factor that is the only gene required to initiate the formation of testis in bipotential gonad (Koopman et al. 1991). On the other hand, in *Drosophila* the Y chromosome plays no role in sex determination. Rather, it is the ratio of X chromosomes to autosomes that determines whether a cell is male or female (Bridges 1916). Determining whether *Hydractinia* follows one of these strategies or an alternative one will require further study. Future cytogenetic analysis of male and female colonies should also reveal whether *Hydractinia* has dimorphic sex chromosomes.

Another question is when and where sex determination occurs in *Hydractinia. Hydractinia* colonies continue to grow new gonozooids throughout their lives. Within each, i-cells continuously differentiate into germline progenitors (Weismann 1883; Müller 1964; DuBuc et al. 2020). One possibility is that the sex determination pathway is activated each time an i-cell commits to gametogenesis. This would likely occur within the germinal zone of the gonozooid after i-cells begin to express *Tfap2*, a transcription factor that commits i-cells to a germ cell fate (DuBuc et al. 2020).

Regardless of the mechanism and location of sex determination, our data indicate the sex determining factor(s) lie within the non-recombining region of the X and Y chromosomes. Although we identified several candidate genes, this list is probably incomplete. One reason is that the sex chromosomes are likely to include at least some of the 4,265 genes that could not be placed in our linkage map (Figure 1F). In addition, the estimated physical size of the *Hydractinia* genome is 514 Mb (Frank et al. 2020), but the genome assembly is only 400 Mb. While most unassembled sequences are probably repetitive elements, some may include protein coding genes or important regulatory elements. This may be exacerbated in regions like Y chromosomes, which often accumulate repetitive elements (Bachtrog 2013; Śliwińska et al. 2016). Finally, because the reference genome was assembled from a male colony, each contig in linkage group 4 could represent an X-specific DNA sequence, a Y-specific DNA sequence, the consensus sequence of a region conserved between X and Y, or a patchwork of X-and Y-specific regions connected by conserved sequences. With these caveats in mind, our analysis of the *Hydractinia* sex locus should be viewed as a ‘first look’ to inspire future studies.

One takeaway from this ‘first look’ is that the sex locus does not appear to contain any of the so-called ‘usual suspects’ for metazoan sex determination genes (Herpin and Schartl 2015). These include Sox family transcription factors and parts of the TGF-beta signaling pathway in vertebrates, *transformer* genes in insects, and *feminizer* genes in a variety of invertebrates. It also includes ‘Doublesex and Mab-3’ (DM) domain-containing genes, which have been found in the sex differentiation pathways of most species studied to date (Herpin and Schartl 2015; Picard et al. 2015). This raises the question of whether the *Hydractinia* sex determination pathway is homologous to the pathways in other animals. Thus, it is germane to note that DM domain-containing genes have been identified in most animal genomes, including cnidarians (Wexler et al. 2014). In addition, at least one DM domain-containing gene has male gonozooid-specific expression, although its function has not been determined (DuBuc et al. 2020). Thus, *Hydractinia* might follow a familiar evolutionary pattern where the overall sex determination and differentiation pathways are conserved and a gene from that pathway has been coopted as the primary sex determination signal (Bachtrog et al. 2014). Alternatively, the primary sex determination signal could be a novel one. This would not be unprecedented; in salmonids, the master sex determining gene *sdY* evolved via duplication from *interferon regulatory factor 9*, an immunity-related gene with no known function in gonad development (Yano et al. 2012, 2013).

One perplexing finding was the presence of two colonies that were initially classified as male but later observed with male and female gonozooids. Male-and female-specific explants from these chimeras have remained exclusively one sex for more than three years. The direction of this sexual chimerism (male to female) is different from that which has been reported previously (Bunting 1894; Hauenschild 1954; Mali et al. 2011). Together, these observations suggest that, although *Hydractinia* has genetic sex determination, abberations may occur that can cause gonads to produce both eggs and sperm or colonies to produce both male and female gonozooids. An alternative explanation in the case of our study is that each sexual chimera was the product of an undetected fusion between juvenile male and female colonieswhere i-cells from both sexual identities persisted in the chimera, but the male matured sooner than the female.

In summary, we have shown that *Hydractinia* has XY sex determination and identified a ∼6.5 Mb region that encodes many promising candidates for the primary sex determiation signal. This work, which maps a sex locus in a genetically tractable cnidarian is a major step toward elucidating sex determination pathways in a broader diversity of animals. With these tools in hand, interested scientists should be able to generate data to bring to bear on questions regarding the sex determination outside of bilaterians in during early animal evolution.

## MATERIALS AND METHODS

### Animal Maintenance and Generation of the Mapping Population

Colonies were maintained and bred at the University of Pittsburgh as previously described (Sanders et al. 2018). The mapping population was created by crossing a male colony (291-10) to a female colony (295-8). Offspring were observed weekly until their sex could be determined. Details are described in the *SI Appendix*.

### Sequencing and Genotyping

DNA was extracted from colonies as detailed in the *SI Appendix*. Illumina libraries were constructed and paired-end sequencing performed at the NIH Intramural Sequencing Center. DNA sequences for the male parent were obtained by downsampling a dataset of Illumina short-read sequences (See *SI Appendix*). Raw reads were mapped to an assembly of the paternal genome with BWA-MEM (Li 2013), duplicates removed with Picard (2019), and genotypes called with GATK HaplotypeCaller (McKenna et al. 2010). Variants suitable for pseudotestcross analysis were then obtained by quality filtering the resulting dataset according to GATK best practices and additional criteria. See *SI Appendix* and https://github.com/nicotralab/chen-et-al-sex-determination (Chen et al. 2022a) for a detailed description of the entire pipeline.

### Genetic Map Construction and QTL Mapping

Genetic maps were constructed in R (version 3.6.1) (R Core Team 2019) with package OneMap (Version 2.1.1) (Margarido et al. 2007). Loci linked to sex were identiried using the R package qtl (version 1.44-9) (Broman et al. 2003). Detailed methods are provided in *SI Appendix* and https://github.com/nicotralab/chen-et-al-sex-determination *(Chen et al. 2022a)*.

### Gene Annotation and Expression

Genes within the non-recombining region of the X and Y chromosomes were identified from annotations of the paternal genome (ref). Gene expression was calculated by mapping sex-specific RNA-seq data (Sanders et al. 2014) to the paternal genome with HISAT2 (Kim et al. 2019) and calculating expression values with Cufflinks (Trapnell et al. 2010). Detailed methods are provided in the *SI Appendix*.

## Acknowledgments

We thank Uri Frank for helpful discussion and Antonio Augusto Franco Garcia and Chris Taniguti for advice on linkage mapping with OneMap. This research is supported by NSF Grant 1923259 (to C.E.S and M.L.N.) by the Intramural Research Program of the National Human Genome Research Institute, National Institutes of Health to A.D.B. (ZIA HG000140). S.M.S. acknowledges support from NIH Grant T32-AI074490.

## Data Availability

Sequence data for colony 291-10 is deposited under BioProject PRJNA807936. Sequence data for colony 295-8 and all F1 offspring is deposited under BioProject PRJNA816479. Additional datafiles and scripts used in our analysis are available at https://github.com/nicotralab/chen-et-al-sex-determination (Chen et al. 2022a) and https://zenodo.org/record/6368105 (Chen et al. 2022b).

