## Supplemental Methods, Figures, and Tables for "XY sex determination in a cnidarian"

**Supplementary Information for**
### XY sex determination in a cnidarian

Ruoxu Chen[1,2], Steven M. Sanders[3,4], Zhiwei Ma[3,4], Justin Paschall[5], E. Sally Chang[5], Brooke M. Riscoe[3,4], Christine E. Schnitzler[6], Andreas D. Baxevanis[5], and Matthew L. Nicotra[3,4,7,*]

[1]School of Medicine, Tsinghua University, Beijing, China

[2]Visiting Scholar, School of Medicine, University of Pittsburgh, Pittsburgh, PA, USA

[3]Starzl Transplantation Institute, Department of Surgery, University of Pittsburgh, Pittsburgh, PA, USA

[4]Center for Evolutionary Biology and Medicine, University of Pittsburgh, Pittsburgh, PA

[5]Computational and Statistical Genomics Branch, National Human Genome Research Institute, National Institutes of Health, Bethesda, MD, USA

[6]Whitney Laboratory for Marine Bioscience, University of Florida, St. Augustine, FL, USA

[7]Department of Immunology, University of Pittsburgh, Pittsburgh, PA, USA

*Corresponding Author: Matthew L. Nicotra

**This PDF file includes:**

Supplementary text
Figures S1 to S5
Tables S1 to S2
Legends for Datasets S1 to S9

**Other supplementary materials for this manuscript include the following:**

Datasets S1 to S9

#### Supplementary Information Text

#### Detailed Materials and Methods

##### Breeding and Animal Maintenance

Colonies were maintained at the University of Pittsburgh as described in (Sanders et al. 2018). Briefly, they were grown on 75 mm x 25 mm glass slides in 38 liter aquaria filled with artificial seawater (Instant Ocean Reef Crystals) and maintained at 22–23 °C. Adult colonies were either fed 3-day-old Artemia nauplii (3x per week) or a suspension of pureed oysters (2x per week). Breeding colonies were kept on a 8h/16h light/dark cycle. After their first exposure to light, males and females were placed in separate 3 liter bins, where they released gametes 1–1.5 h later. Within 20 min of spawning, eggs were harvested by filtering water from the female bin through a 20 µm cell strainer, and sperm was harvested by collecting 10–15 ml from the male container. Eggs were fertilized by mixing them with the sperm and 15 ml additional artificial seawater in a 100 mm polystyrene petri dish. Embryos developed into larvae and were settled 72–96 h post fertilization (hpf). Metamorphosis was induced by incubating the larvae in 56 mM CsCl in filtered seawater for 4–6 h, then transferring them with glass pipettes onto glass microscope slides. Three days later, larvae that successfully metamorphosed began feeding as described above, but without the oyster supplement.

##### Mapping population and sample preparation

A mapping population was created by crossing a male colony (291-10) to a female half-sibling (295-8). The resulting $F_1$ offspring were observed weekly until male or female gonophores could be discerned, at which point they were scored appropriately and moved to male-only or female-only tanks. Thereafter, colonies were observed and cleaned biweekly.

DNA was extracted from colonies when they had grown to cover approximately 2 $cm^2$ of their slide. In most cases, this was performed before the animals had been classified as male or female. Animals were starved for at least two days, then a portion of the colony measuring ~1 $cm^2$ was removed by scraping it from the slide with a razor blade. Harvested tissue was placed in a 1.7 mL microfuge tube, briefly spun in a benchtop microcentrifuge, and residual seawater removed by aspirating with a pipette. Tissue was lysed by the addition of 200 µL UEB buffer (7 M Urea, 0.3125 M NaCl, 0.05 M Tris-HCl pH 8.0, 0.02 M EDTA pH 8.0, 1% N-Lauroylsarcosine sodium salt) (Chen and Dellaporta 1994), followed by grinding with a plastic pestle until all tissue was dissolved. Next, one volume of equilibrated phenol:chloroform:isoamyl alcohol (25:24:1) was added and the mixture homogenized by inverting vigorously. The mixture was centrifuged for 10 minutes at >3000g, and the aqueous phase transferred to a new tube. Total nucleic acid was precipitated with 0.7 volumes isopropanol, then centrifuged for 30 minutes at top speed (at least 13,000 rpm) at room temperature. The resulting DNA pellet was washed twice with 70% ethanol, then transferred to a new tube, centrifuged for 1 minute, and excess 70% ethanol aspirated with a pipette. To remove RNA, the DNA pellet was resuspended in 1X TE, and 1 µL Ambion RNase cocktail (ThermoFisher, Cat. AM2286) per 100 µL suspension was added, followed by incubation at 37℃ for 15 minutes. DNA was then re-extracted with phenol:chloroform:isoamyl alcohol (25:24:1), precipitated with isopropyl alcohol, washed with ethanol, and resuspended with 1XTE as described above. DNA samples were stored at -20°C prior to being sent to the NIH Intramural Sequencing Center for sequencing.

**Karyotype analysis**

Embryos from a cross between colony 291-10 and colony 295-8 were used to generate multiple embryos for karyotyping. Embryos at the 64-128 cell stage (approximately 8 hours post-fertilization) were submitted to the University of Pittsburgh Cell Culture and Cytogenetics Facility. A large number of these embryos were split into three tubes with 3 ml of seawater each. Two tubes were treated with 60µl ColcemidTM (0.1 µg/mL) and one tube was treated with 60µl vinblastine (0.01 mg/ml) (f.c. 0.2ng/µl) for 90 minutes with gentle shaking. Following mitotic arrest, one Colcemid tube was treated with 5 ml 1:1 (sterile water:Mg+Ca+ free seawater) hypotonic solution with 50µl Colcemid for 30 min with constant agitation; the vinblastine tube was treated with 5 ml 1:1 (sterile water:Mg+Ca+-free seawater) hypotonic solution with 50µl vinblastine for 30 min with constant agitation, and the third tube with 90 min Colcemid was treated with 0.075M KCl with 50µl Colcemid for 30 minutes. All of the tubes were fixed with Carnoy's fixative and stored at -20°C overnight. Slides were prepared the next day following a few washes in the fixative. Both Colcemid-treated samples produced a few metaphase cells, but the vinblastine tube produced no visible metaphase cells. All of the slides were observed on an Olympus BX61 microscope and imaged and analyzed using the Genus software platform on the Cytovision System (Leica Microsystems, San Jose, CA). Nineteen out of twenty cells examined expressed a near-diploid chromosome complement with chromosome numbers of 30 per cell (2n=30). One cell had a near-tetraploid chromosome number with an approximated 54 chromosomes. An attempt to karyotype three cells revealed what appeared to be a normal diploid chromosome complement with 15 pairs of chromosomes (2n=30) in the poorly G-banded metaphase cells.

**Whole genome sequencing and SNP discovery**

A detailed description of the complete analysis pipeline, including all scripts downloaded from https://github.com/nicotralab/chen-et-al-sex-determination (Chen et al. 2022a). Several data sets were too large to be placed in the GitHub repository but can be downloaded directly from https://zenodo.org/record/6368105 (Chen et al. 2022b).

For the female parent and all $F_1$ progeny, PCR-free libraries were generated from 1 µg genomic DNA using the TruSeq® DNA PCR-Free HT Sample Preparation Kit (Illumina). The median insert sizes were approximately 400 bp. Libraries were tagged with unique dual index DNA barcodes to allow pooling of libraries and minimize the impact of barcode hopping. Libraries were pooled for sequencing on the NovaSeq 6000 (Illumina) to obtain at least 500 million 151-base read pairs per individual library. All raw sequence data for the female parent and offspring are available via BioProject PRJNA816479 at NCBI. The male was previously sequenced to generate 240 million reads (BioProject: PRJNA807936). For this study, the male data files were downsampled with seqtk (Li 2019), resulting in 66 x $10^6$ 251 bp paired end reads. The downsampled files can be downloaded from https://zenodo.org/record/6368105 (Chen et al. 2022b)

For each sample, raw reads were mapped to an assembly of the paternal genome, which can be downloaded at https://research.nhgri.nih.gov/hydractinia/download/assembly/symbio/Hsym_primary_v1.0.fa.gz. Reads were mapped to the assembly using BWA-MEM (Li 2013) with mapping parameters '-M -t 8'. The resulting .sam files were converted to .bam format and then sorted with samtools (Li et al. 2009). Duplicates were then marked with Picard (Picard toolkit 2019) . Genotypes were called with GATK HaplotypeCaller (McKenna et al. 2010). The resulting file, rawvariants.90f1.vcf.gz, can be downloaded from https://zenodo.org/record/6368105 (Chen et al. 2022b)

Raw variant calls were filtered to generate datasets of high quality variants suitable for genetic mapping. Briefly, raw variant calls were filtered with a custom python script (qualityfilter.py) to retain only those variants for which (1) no samples were missing data; (2) all samples genotyped as homozygous reference (0/0) had no more than two mapped reads corresponding to the alternative allele and also had more than ten mapped reads corresponding to the reference; (3) all samples genotyped as homozygous alternative (1/1) had no more than two mapped reads corresponding to the reference allele and also had more than ten mapped reads corresponding to the alternate allele; 4) all samples genotyped as heterozygous (0/1) had an alternate allele read count percentage of greater than 0.3 or less than 0.7. This dataset was further filtered according to GATK best practices (Van der Auwera et al. 2013). Specifically, SNPs were flagged as low quality if they met any of the following criteria: quality by depth (QD) < 2; Fisher's Exact Test of strand bias (FS) > 60; RMS mapping quality (MQ) < 40; rank sum of alt versus reference mapping quality (MQRankSum) < 12.5; read position rank sum (ReadPosRankSum) < 8; and read depth (DP) < 10. Indels were flagged as low quality if they met any of the following critera: QD < 2.0, FS > 200, or ReadPosRankSum < -20.0. Flagged variants were then removed with bcftools (Li et al. 2009). The resulting file, GATK-passed.vcf can be downloaded from https://zenodo.org/record/6368105 (Chen et al. 2022b)

From this filtered dataset, we generated two sets of markers suitable for mapping via a pseudo-testcross strategy (Grattapaglia and Sederoff 1994). In a pseudo-testcross, two parents from an outcrossing population are bred to create an $F_1$ population. A genetic map for each parental genome is then constructed with markers that are heterozygous in that parent and homozygous in the other. For example, a genetic map of the maternal genome can be constructed from variants that are homozygous in the male parent and heterozygous in the female parent (i.e. "0/0" x "0/1" in the notation of a .vcf file). Likewise, a genetic map of the paternal genome can be constructed from variants heterozygous in the male parent and homozygous in the female parent (e.g. "0/1" x "0/0"). To create a set of variants for mapping the maternal genome we used the linux command-line tool "awk" to extract variants with paternal genotype "0/0" and maternal genotype "0/1" (hereafter, the "female PT dataset"). Note that variants with paternal genotype "1/1" were not included because the paternal genome was the reference genome, thus no paternal genotype should be homozygous for the alternative allele. To create a dataset of markers to map the paternal genome we used "awk" to extract variants with paternal genotype "0/1" and maternal genotype "0/0" or "1/1" from the filtered dataset (hereafter, the "male PT dataset").

Next, we used a custom bash script (getPTvariants.sh) to identify probable genotyping errors according to the segregation pattern of offspring genotypes. In both of the resulting datasets, the segregation of $F_1$ genotypes is expected to be 1:1 homozygous:heterozygous. Homozygous offspring should have the same genotype as the homozygous parent (e.g., if one parent is "0/0" and the other parent "0/1", homozygotes should be "0/0"). An $F_1$ offspring with a genotype of the alternative homozygote class ("1/1" in the preceding example) would probably be the result of a genotyping error. We determined the frequencies of such errors for each $F_1$ offspring, and found they had an average of 1.18% ± 0.21% (mean ± standard deviation ) in the female PT dataset and 1.24% ± 0.19% in the male PT dataset.

Unexpected homozygous genotypes could also arise in $F_1$ offspring if one of the two parents were misgenotyped. At such a variant, the unexpected homozygote class should segregate with other genotypes in a mendelian pattern. To search for this type of genotyping error, we determined the frequency of unexpected homozygous genotypes at each variant in both datasets. We found 5.12% of variants in the female PT dataset and 4.9% of variants in the male PT dataset had unexpected homozygotes. At each of these variants, the number of unexpected homozygotes averaged 24.59% ± 0.68% in the female PT dataset and 23.69% ± 1.13% in the male PT dataset. The occurrence of unexpected homozygotes at a frequency of ~0.25 is consistent with both parents being heterozygous at these variants. To exclude these variants, as well as the genotyping errors described above, we changed all unexpected homozygotes to missing data, then excluded variants with more than 10% missing data from each PT dataset. The resulting files, GATKBP-passed.femaleHet.abxabRemoved.vcf and GATKBP-passed.maleHet.abxabRemoved.vcf can be downloaded from https://zenodo.org/record/6368105 (Chen et al. 2022b).

**Genetic Map Construction**

Genetic maps were constructed in R (version 3.6.1) (R Core Team 2019) with package OneMap (Version 2.1.1) (Margarido et al. 2007). Prior to importing the datasets into R/Onemap, variants were tested for Mendelian segregation ($\chi^2$ goodness of fit) and those where $p < 0.00001$ were removed with a custom Perl script (removeDistorted.pl) The resulting files, femalePT.vcf.gz and malePT.vcf.gz are available at https://github.com/nicotralab/chen-et-al-sex-determination (Chen et al. 2022a). Each dataset was then thinned with vcftools (Danecek et al. 2011) to ensure that the distance between adjacent variants was no less than 5,000 bp and converted from vcf format into OneMap's ".raw" format with a bash shell script (thin-for-onemap.v2.sh) to generate Datasets S2 and S3. After importing into OneMap, variants with identical genotypes were binned to a single marker with the function find_bins(). The remaining non-redundant markers were re-tested for segregation distortion at $p < 0.05$, with Bonferroni's multiple test correction applied. Two-point tests were used to calculate recombination fractions and LOD scores for each pair of markers, and linkage groups between non-distorted markers identified with a maximum recombination fraction (rf) of 0.4 and minimum LOD score determined by the OneMap function suggest_lod().

To order markers within each linkage group, the function order_seq() was used. This function selects an initial set of five markers and applies an exhaustive search to determine the order with the lowest LOD score. To this framework map, the remaining markers are added one-by-one to optimize the total LOD score of the growing map. Recombination fractions were converted to centiMorgan (cM) units using the Kosambi map function.

Most of the initial maps contained pairs of markers that were placed within 0.0001 cM of each other by the OneMap software. Upon closer inspection, we discovered that these markers were simply markers that were located on the same contig in the reference genome, but had their alternative alleles in opposite phase of one another. Since these markers were essentially redundant to one another we decided to remove them from the maps. To do this we identified them in the initial maps with a custom perl script (identify_redundant_markers.pl), then removed them using the drop_marker function in OneMap.

A recombination fraction plot was then generated with the function rf_graph_table and visually inspected to identify misplaced markers. These were removed from the map and re-inserted with the try_seq() function. Markers that could not be confidently placed were removed entirely from the final maps. Summary statistics for each map were calculated using the Genetic Map

Comparator (Holtz et al. 2017). The 'unbinned' maps were created by using the custom perl script unbin_markers_in_map.pl.

##### Comparison of recombination rates

To compare recombination rates in the female and male genomes, we identified pairs of markers in the final maps that were located within 5 kb of each other in the reference genome assembly. Linkage groups having two or more such markers were then reconstructed as described for the initial maps. Linkage maps were then compared and summary statistics calculated with the Genetic Map Comparator (Holtz et al. 2017).

##### QTL mapping

Loci linked to sexual phenotype were identified using in R with package qtl (version 1.44-9) (Broman et al. 2003). Prior to QTL mapping, the data were prepared for R/qtl with a custom perl script (onemapRaw_to_Rqtl.pl). Briefly, for each PT dataset, the marker data in OneMap's .raw format and the corresponding linkage map were combined and converted to R/qtl's .csvr format. Header information including the phenotypes (sex) of each colony was added manually. The two resulting files (femaledata.rqtl.with.phenotypes.csvr and maledata.rqtl.with.phenotypes.csvr available in from https://github.com/nicotralab/chen-et-al-sex-determination) were imported into R using the read.cross function. For each dataset, QTL genotype probabilities were calculated with calc.genoprob with default settings and the kosambi map function. A single-QTL genome scan was performed using the function scanone for a binary phenotype under the Haley-Knott regression model, with 1000 permutations and perm.Xsp = 0. Significance thresholds were calculated with the summary function. Figures were created with the plot function, then imported into Adobe Illustrator for further annotation.

##### Gene annotation

Gene functions were predicted using the standalone version of PANNZER2 (Koskinen et al. 2015) with default parameters, as well as with a DIAMOND (Buchfink et al. 2015) similarity search against nr with the option 'blastp'. Homologous sequences in the NCBI Model Organisms (landmark) database were identified using BLASTP as implemented through the BLAST website (https://blast.ncbi.nlm.nih.gov/Blast.cgi). Conserved protein domains were identified with the Pfam database using hmmscan as implemented by the HMMER Webserver (Potter et al. 2018).

##### Gene Expression Analysis

Sex specific expression of genes in the putative sex determination locus was estimated using previously published RNA-seq libraries from *H. symbiolongicarpus* gastroozooids and gonozooids (Sanders et al. 2014) (SRA Accessions SRX871546-9 and SRX871551-4). The paired-end RNA-seq reads from each library were mapped to the entire genome assembly using HISAT2 (Kim et al. 2019) under default settings. The resulting mapping files were processed and sorted using samtools (Li et al. 2009) before proceeding to quantitation. Using the reference annotations for the primary haplotype of the genome, FPKMs of each gene model was estimated for each library and normalized by library size using the cuffnorm function of Cufflinks (Trapnell et al. 2010).

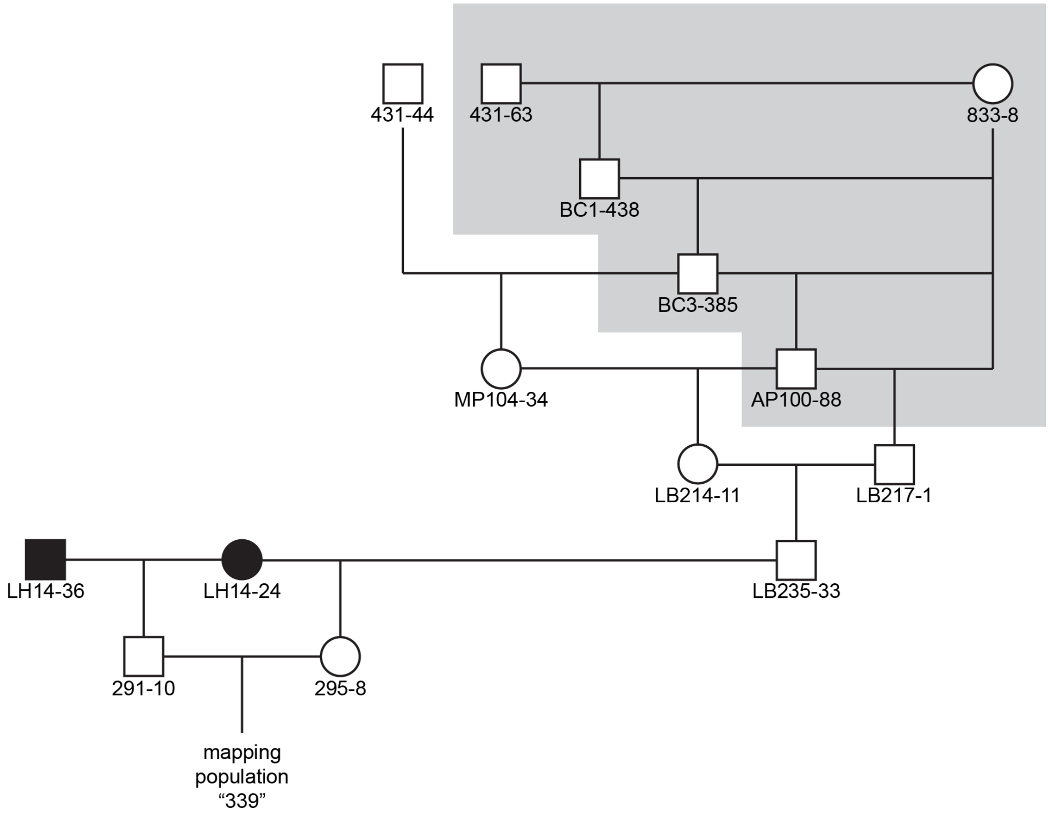


**Fig S1. Pedigree of the colonies used to generate the mapping population.**
Field-collected colonies are denoted with black symbols. Colony 291-10 is the offspring of two colonies collected from Lighthouse Point, New Haven, CT in 2014. Colony 295-8 is the offspring of a field collected colony and a laboratory strain, 235-33. The pedigree of colony 235-33 can be recreated by concatenating previously published pedigrees (shaded area) (Cadavid et al. 2004; Powell et al. 2007). Colony AP100-88 is from the mapping population in Powell et al. (2007). Colony 431-44 is from the mapping population in Cadavid et al. (2004).

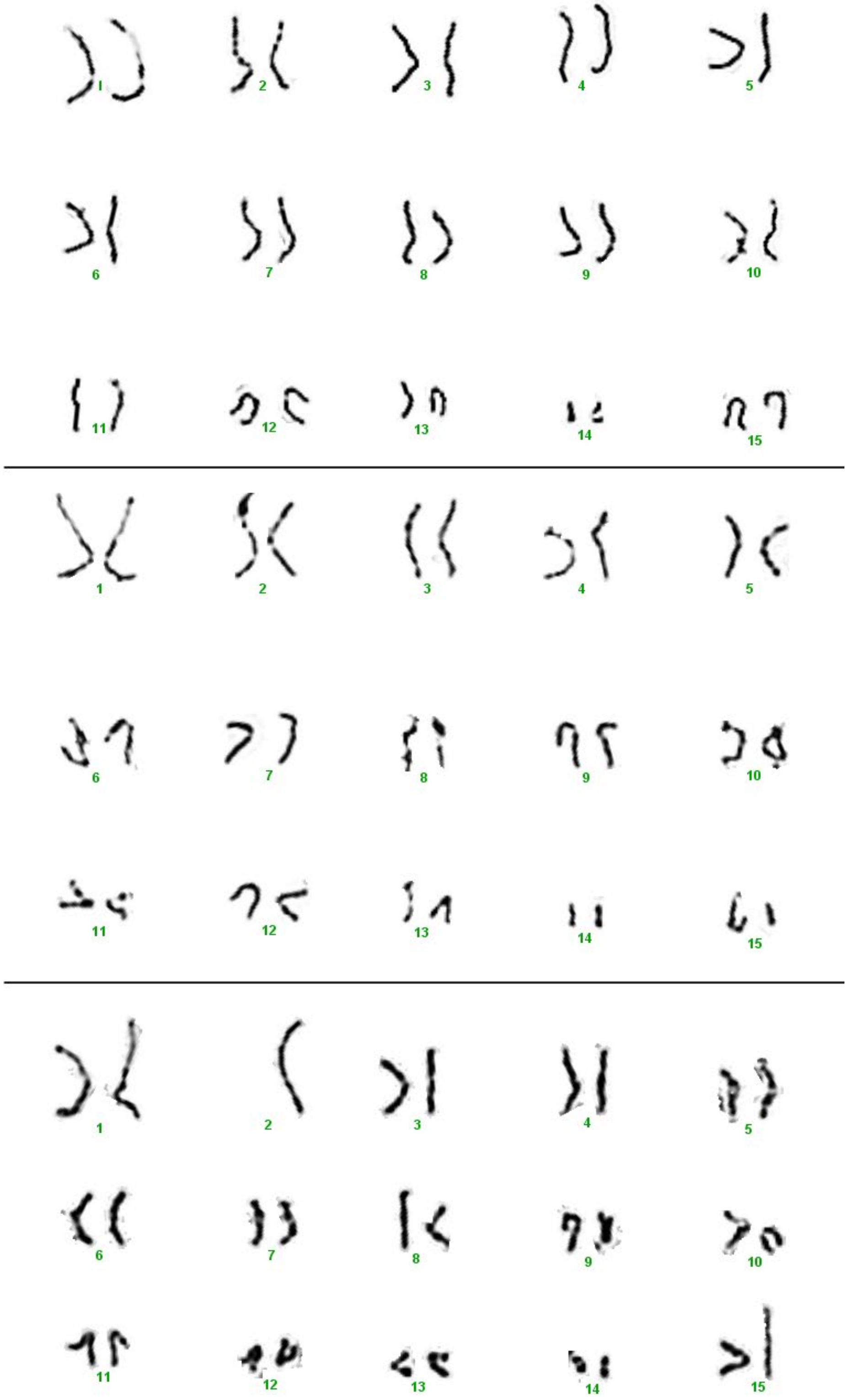


**Fig S2. Karyotype analysis of *Hydractinia symbiolongicarpus***
Karyotype of three metaphase cells from 64-128 cell embryos. Despite poor G-banding, each had what appeared to be a normal diploid chromosome competent of 15 pairs (2n = 30).

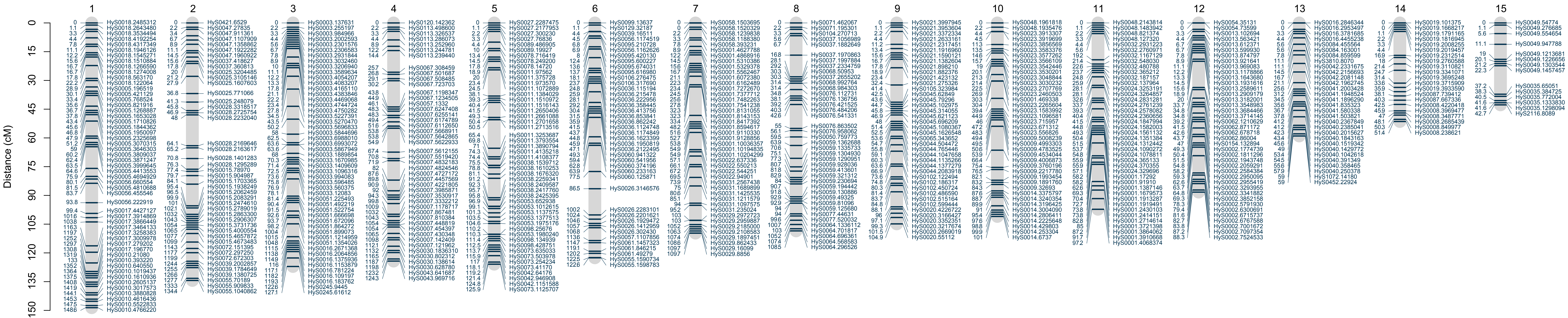


**Fig. S3. Maternal linkage map of the Hydractinia genome.** Marker names (contigname.position) are displayed to the right of each linkage group. Genetic position, in centimorgans, is displayed to the left.

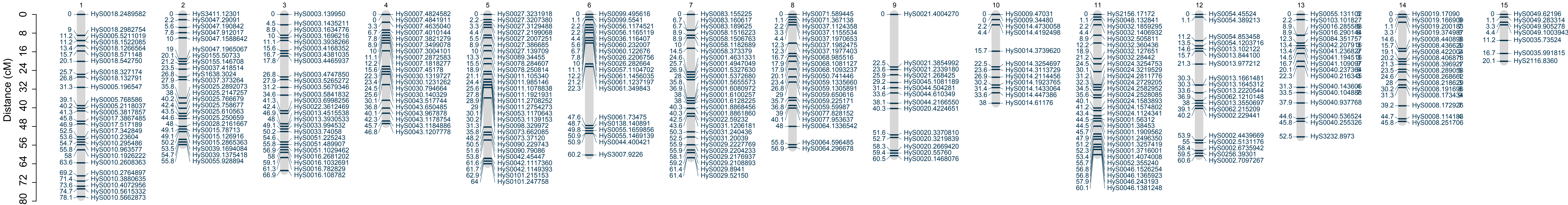


**Fig. S4. Paternal linkage map of the Hydractinia genome.** Marker names (contigname.position) are displayed to the right of each linkage group. Genetic position, in centimorgans, is displayed to the left.

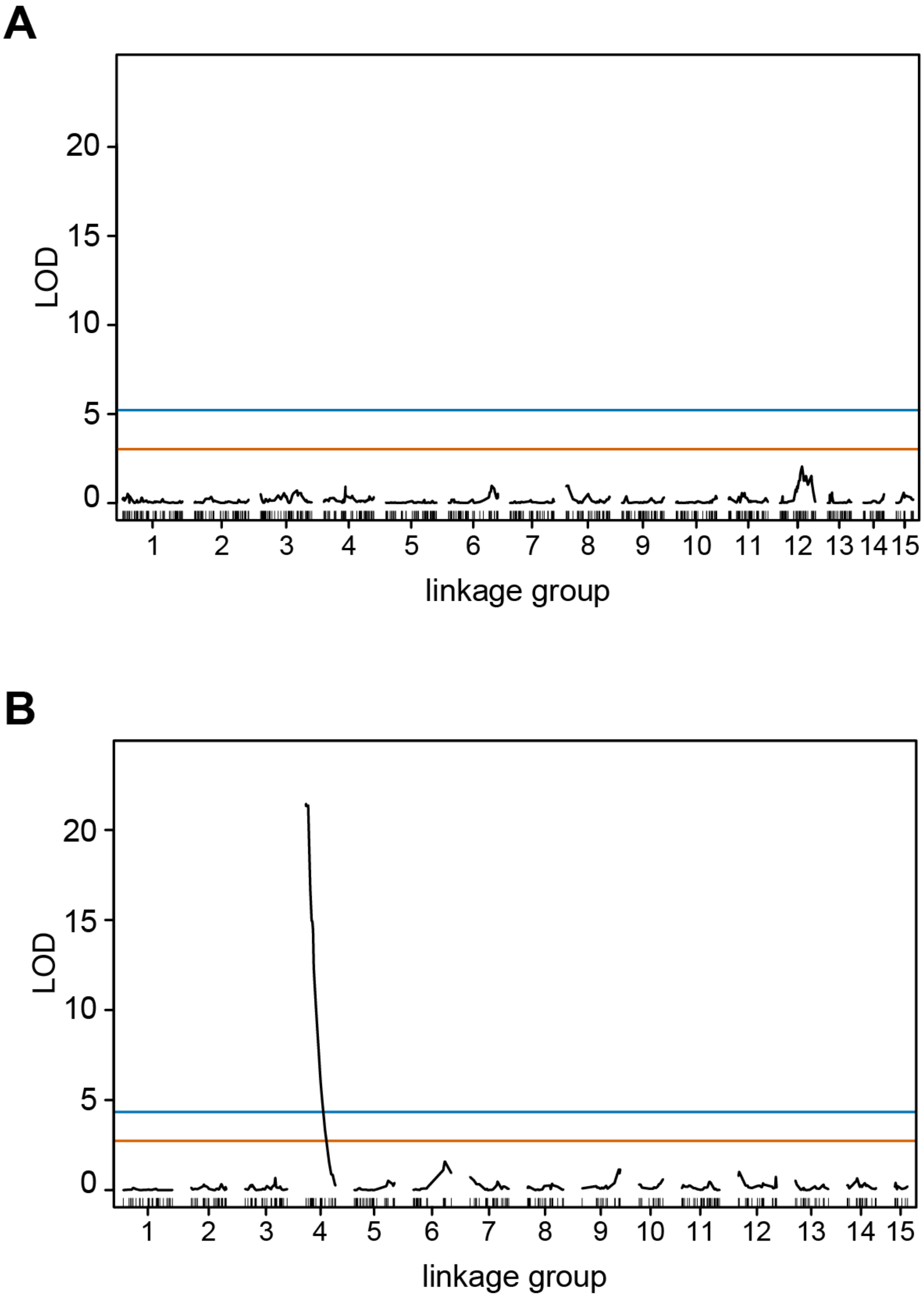


**Fig. S5.** QTL analysis excluding sexual chimeras.
*(A)* LOD chart of QTL for sex in the maternal linkage map. *(B)* LOD chart of QTL for sex in the paternal linkage map. The blue

**Table S1. Representation of the assembled *Hydractinia* genome in linkage maps**

| placement in linkage maps | contigs | | base pairs | | annotated genes | |
|---|---|---|---|---|---|---|
| both | 160 | *(3.5)** | 265,235,647 | *(65.7)* | 17,444 | *(79.2)* |
| maternal only | 72 | *(1.6)* | 3,850,356 | *(1.0)* | 127 | *(0.6)* |
| paternal only | 41 | *(0.9)* | 3,770,032 | *(0.9)* | 186 | *(0.8)* |
| none | 4,237 | *(95.0)* | 130,968,334 | *(32.4)* | 4,265 | *(19.4)* |
| **TOTAL** | **4,510** | | **403,824,369** | | **22,022** | |

* percent of total indicated in parentheses

**Table S2.** Contigs with variants in multiple linkage groups

| Contig | Linkage group in maternal map | Linkage group in paternal map |
|---|---|---|
| HyS0001 | 7, 11 | 7, 11 |
| HyS0013 | 3, 12 | 3, 12 |
| HyS0016 | 3, 13 | 3, 13 |
| HyS0022 | 3, 7 | 3, 7 |
| HyS0030 | 4, 11 | 4, 11 |
| HyS0037 | 2, 8 | 2, 8 |
| HyS0039 | 2, 6, 10 | 2, 10 |
| HyS0042 | 3, 5, 13 | 5, 13 |
| HyS0043 | 2, 4 | 2, 4 |
| HyS0048 | 10, 11 | 10, 11 |
| HyS0050 | 8, 10 | 8, 10 |
| HyS0055 | 2, 6 | 2, 6 |
| HyS0056 | 1, 6 | 1, 6 |
| HyS0057 | 4, 6 | 4, 6 |
| HyS0122 | 10, 14 | 10, 14 |

**Other supplementary materials for this manuscript include the following:**

**Table S3** (separate file). A tab delimited text file with gene expression data and functional annotations for each gene in the non-recombining region of the X and Y chromosomes.

**Dataset S1** (separate file). A tab delimited text file with per sample sequencing statistics.

**Dataset S2** (separate file). A space-delimited text file of genotype data in R/Onemaps ".raw" format representing the female PT dataset.

**Dataset S3** (separate file). A space-delimited text file of genotype data in R/Onemaps ".raw" format representing the male PT dataset.

**Dataset S4** (separate file). A space-delimited text file representing the maternal linkage map. Columns denote linkage group, marker name, and genetic position in centimorgans.

**Dataset S5** (separate file). A space-delimited text file representing the paternal linkage map. Columns denote linkage group, marker name, and genetic position in centimorgans.

**Dataset S6** (separate file). A space-delimited text file showing all markers from Dataset S2 that can be placed on the maternal linkage map.

**Dataset S7** (separate file). A space-delimited text file showing all markers from Dataset S3 that can be placed on the paternal linkage map.

**Dataset S8** (separate file). A text file containing FASTA formatted nucleotide sequences of 461 genes located in the non-recombining region of the X and Y chromosomes.

**Dataset S9** (separate file). A tab-delimited text file with gene expression information and annotations for each gene in the non-recombining region of the X and Y chromosomes. Each field is described in the comments at the beginning of the file.
